# Metabolic Reprogramming: Short-term Western Diet Exposure Induces Sustained Changes in Plasma Metabolites

**DOI:** 10.1101/554980

**Authors:** Biswapriya B. Misra, Ram P. Upadhayay, Vicki Mattern, John S. Parks, Laura A. Cox, Anthony G. Comuzzie, Michael Olivier

## Abstract

Dietary patterns are well known to contribute to the metabolic syndrome and related disorders. We evaluated the impact of a short-term Westernized diet (WD) exposure on systemic plasma metabolomic changes in a cohort of adult male baboons. In this pilot study, five male baboons (n=5) raised and maintained on a standard monkey chow diet (high complex carbohydrates, low fat) were exposed to a challenge WD (high in saturated fats and simple carbohydrates) for 7 weeks (49 d), followed by a 57-day washout period on monkey chow (106 d). In addition to monitoring clinical measures, we used a comprehensive two-dimensional gas chromatography-time-of-flight mass spectrometry (2D GC-ToF-MS) platform to assess the metabolomic changes in plasma at three time points. Twenty-three metabolites were changed in response to the WD (49 d), but the response across the animals was highly variable. All animals presented a very different metabolic profile than at baseline (0 d), and the washout period resulted in a relatively homogenous metabolic state across all animals. A short-term exposure to a WD led to long-term changes in the metabolic profile, suggesting a “reset” of the metabolic program.

## 1. Introduction

In 2015, an estimated 415 million adults were living with diabetes, and 5 million deaths were attributable to the disorder. The total global health expenditure due to diabetes was estimated at 673 billion USD, and predicted to rise significantly by 2040 [1]. Similar trends have been reported for obesity, with 107.7 million children and 603.7 million adults obese in 2015, and 4 million deaths globally attributed to obesity [2]. Changes in nutrition and lifestyle over the past decades have likely significantly contributed to this unprecedented rise in obesity and diabetes rates. While dietary saturated fats are recognized as a significant contributor to impaired cardiometabolic health, dietary fats alone do not seem to account for the total metabolic disruption associated with increasing risk of cardiovascular disease (CVD) and other comorbidities such as type 2 diabetes (T2D). There is emerging evidence that simple carbohydrates in the diet also negatively impact overall cardiometabolic health in conjunction with saturated fats, and that this combination of dietary elements gives rise to the full spectrum of metabolic disruption and dysregulation (elevated plasma triglyceride levels, decreased insulin sensitivity, and increasing fat accumulation). In 2009, the American Heart Association published a statement on the negative impact of a diet high in simple carbohydrates on overall cardiovascular health [3]. Unfortunately, from a public health perspective, the vast majority of individuals who typically consume a diet high in saturated fats also tend to consume higher amounts of simple carbohydrates, and this combination has been termed a Western diet (WD). A higher Western dietary pattern score is associated with an increased risk for type 2 diabetes, coronary heart disease, and colon cancer [4-8]. While dietary saturated fats and simple carbohydrates separately negatively impact cardiometabolic health, these two dietary elements consumed together likely act synergistically to increase both the rate of disease progression as well as its severity.

Long-term changes in body weight correlate with characteristic changes in serum metabolite concentrations [9]. Changes in plasma triglyceride, cholesterol, and glucose levels are often clinical manifestations of weight gain in response to a WD. Interestingly, improved nutrition, dieting and weight loss do not invariably lead to a return to normal plasma metabolite concentrations, and these altered plasma levels of specific metabolites may be contributing to the increased cardiovascular disease risk observed in patients with repeated cycles of weight gain and loss (often termed yoyo dieting or weight cycling) in both females [10, 11] and males [12]. However, the exact pattern of metabolic changes during this weight cycling has not been characterized, and it is unknown how quickly a WD exposure (with or without weight gain and subsequent loss) causes long-term changes.

With the advent of modern metabolomics technologies, such as gas chromatography-coupled mass spectrometry, it is now possible to conduct comprehensive and unbiased analyses of serum samples to characterize the complex metabolic changes resulting from weight gain and changes in diet patterns. In a recent study, an 8% weight increase was induced within 6-8 weeks in human volunteer subjects on a WD [13]. A detailed metabolomics analysis revealed significant changes in response to the diet change and weight gain in study participants. The analysis suggested that even short-term administration of a WD, and the resulting short-term weight gain, led to metabolomic changes that persisted even after participants were returned to a normal diet, and the initial weight gain was reversed. However, the findings were highly variable among individuals, suggesting inter-individual factors (such as genetic factors or past life style) also contributing to the observed response and highlighting the challenges to investigating this phenomenon in adult human patients or volunteers.

Non-human primate (NHP) models, such as baboons, have been used extensively to study cardiometabolic disorders in response to Western and other diets [14-17]. These studies benefit from the similarity in metabolism and nutritional needs of these animals compared to humans, and a controlled environment in which diets of defined composition can be fed without having to account for variable prior exposures of animals.

For this study, we formulated a WD, mimicking the Westernized diet consumption in humans that is high in both saturated fats and simple carbohydrates. Healthy adult baboons (*Papio hamadryas*) were exposed to this challenge diet for 7 weeks, and then returned to normal monkey chow, a healthy low fat complex carbohydrate diet. We applied metabolomic analyses to evaluate the changes induced by this WD exposure, as reflected in the composition and distribution of plasma metabolites, and then assessed whether a return to a normal diet reversed the WD-induced changes. As we describe below, we observed significant changes in plasma metabolites in baboons after 7 weeks on the WD, independent of weight gain or other clinical serum measures. Interestingly, metabolite levels did not return to pre-challenge levels even after 56 d of monkey chow consumption, suggesting that even a short-term exposure to a WD leads to long-term alterations of basic metabolite levels.

## 2. Experimental Section

### 2.1 Baboon study cohort and composition of challenge diet

For this study, we selected five adult male baboons (*Papio hamadryas*) maintained as part of the baboon colony at the Southwest National Primate Research Center (Texas Biomedical Research Institute, San Antonio, TX). All animals were 7 years of age and had an average weight of 29.4 kg (26.6 – 32.5 kg). All animals were raised and maintained on a standard monkey chow diet (high complex carbohydrates, low fat) prior to the start of this dietary challenge. Each animal was exposed to the WD (high in saturated fat and simple carbohydrates) for a period of 49 d, followed by a 57 d washout period (106 d) where the baboons were returned to the baseline monkey chow diet. The caloric content of the monkey chow and WD were 3.25, and 4.59, Kcal/g respectively. The protein content of the chow and WDs were held constant between the diets while the concentrations and types of carbohydrates were modified. Further, in the WD, 40% of calories were derived from lard, and simple carbohydrates were predominantly derived from high fructose corn syrup. The detailed diet compositions are listed in **Table 1**. All animals were allowed to feed *ad libitum* throughout the study, without measuring individual food intake. All animal procedures were reviewed and approved by the Texas Biomedical Research Institute’s Institutional Animal Care and Use Committee and performed by veterinary staff at the Southwest National Primate Research Center.

**Table 1.** Comparison ofstudy diets.

|  | Monkey<br>Chow | Western Diet<br>(WD) |
| --- | --- | --- |
| Energy (Kcal/g) |  |  |
|  | 3.25 | 4.3 |
| Carbohydrates (g/100g) |  |  |
| **Total** | 62 | 51 |
| Simple | 3 | 22 |
| Complex | 59 | 29 |
| Protein (g/100g) |  |  |
| **Total** | 28 | 10 |
| Fat (g/100g) |  |  |
| **Total** | 10 | 39 |
| Saturated | 2 | 18 |
| Monounsaturated | 3 | 16 |
| Polyunsaturated | 5 | 5 |
| Trans | 0 | 0 |

### 2.2 Clinical assays

Serum samples collected after overnight fast at baseline (day 0), after 7 weeks on the WD (day 49), and after a washout period on chow diet (day 106) were stored at −80 °C, and used for metabolomics assays as well as standard clinical chemistry measurements. Circulating biomarkers reflecting the status of lipid and glucose metabolism, as well as liver function and general systemic inflammation markers were measured using commercially available human assay kits with protocols established at the Southwest National Primate Research Center for non-human primates. Total serum cholesterol (TSC) and triglyceride (TG) concentrations were determined enzymatically using commercial reagents in a clinical chemistry analyzer. High-density lipoprotein cholesterol (HDLC) was measured in the supernatant after heparin-Mn^+2^ precipitation. Blood glucose was measured using the Alfa Wasserman ACE clinical chemistry instrument (West Caldwell., NJ). Plasma insulin and C-peptide were measured by immunoassay using the Immulite Analyzer (Diagnostic Products Corporation, Los Angeles, CA). Interleukins, IL-6, IL-8 (pg/mL) were assayed using a sandwich-style enzyme-linked immunoassay kit, as suggested by the manufacturer (R&D Systems, Inc., Minneapolis, MN, USA). A high-sensitivity assay kit (Kamiya Biomedical, Seattle, WA, USA), with a latex particle-enhanced immunoturbidometric method, to measure C-reactive protein (CRP) concentrations. Summaries of raw data are presented as means, and standard deviations. A list of all clinical measurements used in this study is provided in **Table S1**. Numerous published studies from the institution have shown the accuracy/precision, sensitivity/specificity of these assays over the years in baboon models and that measures for each are comparable to humans [18-20].

### 2.3 Metabolomic analysis using GC-MS

Gas chromatography-mass spectrometry (GC-MS) was performed as described elsewhere [21, 22] using a two-dimensional gas chromatography time-of-flight mass-spectrometer (2D GC-ToF-MS) platform. Plasma samples (30 μL) were extracted serially, once each using mixtures of isopropanol: acetonitrile: water (3:3:2, v/v/v) and acetonitrile: water (1:1, v/v) following a standard polar metabolite extraction protocol described [23]. Pooled metabolite extracts (1500 µL) from both phases were dried under vacuum at room temperature, and were then sequentially derivatized using methoxyamine hydrochloride (MeOX) and *N*-methyl-*N*-trimethylsilyl-trifluoroacetamide (MSTFA) [21]. Briefly, chemical derivatization was carried out by adding 10 μL of MeOX (20 mgmL^-1^in pyridine) and shaking at 55 °C for 60 min., which was followed by trimethylsilylation reactions at 37 °C for 60 min. after addition of 90 μL of MSTFA. Derivatized samples (1 μL) were injected in splitless mode using an autosampler (VCTS, Gerstel(tm), Linthicum, MD, USA) into the 2D GC-MS system consisting of an Agilent^©^7890 B gas chromatograph (GC) (Agilent Technologies, Palo Alto, CA, USA) connected to a Pegasus ® 4D ToF-MS mass spectrometer (LECO Corp., St. Joseph, MI, USA) equipped with an electron impact (EI) ionization source (70 eV). Injection temperature was maintained at 250 °C, helium was used as a carrier gas with a flow rate of 1 mL min^-1^. Separation on the GC was performed using a primary Rxi®-5Sil MS capillary column (Cat. No. 13623-6850, Restek, Bellefonte, PA, USA) (30 m × 0.25 mm × 0.25 μm) and a secondary Rtx®-5 capillary column (Cat. No. 40201-6850, Restek, Bellefonte, PA, USA) (2 m × 0.15 mm × 0.15 µm). The temperature program was as follows: isothermal at 70 °C for 1 min, 6 °C min^-1^ramp to 310 °C, and a final 11 min hold at 310 °C. The system was left at 70 °C for 5 min before the next injection to allow temperature equilibration. Mass spectra were collected at 200 scans/s over a range of *m/z* 40-600. Transfer line and the ion source temperatures were both set to 280 °C. Pooled samples that served as quality control (QCs) were injected at scheduled intervals for instrumental performance, tentative identification, and monitoring of shifts in retention indices (RI).

### 2.4. Processing of raw GC-MS data

GC-MS data were pre-processed, cleaned, and aligned using ChromaToF version 4.71.0.0 (LECO Corp., St. Joseph, MI, USA) following settings as described elsewhere [22, 24]: viz. S/N: 25; peak width: 0.15, base line offset: 1; m/z range: 40-600. Metabolite identification included consulting the open source MSRI spectral libraries of the Golm Metabolome Database available from the Max-Planck-Institute for Plant Physiology, Golm, Germany (http://csbdb.mpimp-golm.mpg.de/csbdb/gmd/gmd.html) for matching the mass spectra and retention indices (RIs) [25] and the NIST Mass Spectral Reference Library (NIST11/2011; National Institute of Standards and Technology, USA). Base peak areas for individual metabolite spectra were used for quantification. The filtered raw GC-MS data included data (base peak areas) from five biological replicates. Individual metabolite quantification data were assembled into a joint data set which included hits from both the databases.

### 2.5 Statistical analyses

Standard statistical analyses (summary statistics) were performed using the statistical software R (Version 3.5.0, http://www.R-project.org) [2, 26] and Microsoft Excel. Normalized (internal standards), transformed (log_2_), imputed, and scaled peak areas representative of relative metabolite amounts were obtained using DeviumWeb [27] and are presented in tables and figures. Values reported in all tables and text are presented as means, and differences were considered significant when P < 0.05 (nominal P-values). Short Time series Expression Miner (STEM) [28] was used as a Java implementation with a graphical user interface available at http://www.cs.cmu.edu/∼jernst/st/ for clustering the metabolite accumulation patterns according to three time points.

### 2.5.1 Univariate analysis

ANalysis Of VAriance (ANOVA) was performed using R packages. Hierarchical clustering analysis (HCA) using average linkage clustering was performed on Pearson distances using PermutMatrix [29] from the metabolite abundance data. For heat maps displaying the top 25 metabolites, data were normalized using z-scores of the abundances for individual metabolites under the peak area.

### 2.5.2 Multivariate analysis

Principal components analysis (PCA) was performed using the package DeviumWeb [27], where the output consisted of score plots to visualize the contrast between different samples and loading plots to explain the cluster separation. Data were scaled with unit variance without any transformation. Partial least-squares discriminant analysis (PLS-DA) was used to highlight differences between the metabolic phenotypes at three time points (0 d, 49 d, 106 d) in the study.

### 2.6. Pathway enrichment and clustering analysis

Pathway enrichment analysis was performed at MetaboAnalyst (www.Metaboanalyst.ca) [30], and Chemical Translation Service (CTS: http://cts.fiehnlab.ucdavis.edu/conversion/batch) was used to convert the common chemical names into their KEGG, HMDB, CAS, PubChem CID, LipidMAPS IDs and InChiKeys.

## 3. Results and Discussion

### 3.1. Changes in clinical measures in response to short-term WD exposure

We used baboons fed regular monkey chow and a WD as a NHP model to mimic human physiological and metabolic adaptations to dietary fat and sugar overload. The WD diet constituted 7-fold greater simple sugars and 2-fold lower complex carbohydrates, 4-fold higher total fat, 9- and 5-fold higher saturated and monounsaturated fats than the chow diet, mimicking a typical human WD, with only a modest increase in total energy content (41% higher Kcal/g in the WD).

Clinical assays included quantification of glucose, insulin, total cholesterol, LDLC, HDLC, Hb, HbA1c, and triglyceride levels, alongside several cytokines, for the five baboons at baseline and after 49 d of WD challenge. None of the clinical measures were significantly different across all animals at 49 d, but insulin [0.23 fold less at 49 d (260.66 ± 88.41 pg/ml) compared to 0 d (1129.2 ± 470.82 pg/ml)], adropin [4.57 fold increase at 49 d (7.52 ± 9.9 ng/ml)], and interleukin-6 (IL-6) (0.54 fold decrease at 49 d (0.75 ± 0.44 pg/ml)] were dysregulated after 49 d of WD administration, and showed consistent trends across all animals. Adropin is a peptide hormone regulating carbohydrate and lipid metabolism, and implicated in metabolic diseases, central nervous system function, endothelial function and cardiovascular disease that links dietary macronutrient intake with energy homeostasis and lipid metabolism [31][32]. IL-6 is known to be induced by exposure to a high fat diet [33]. Though uncommon, but in one study, when 8 healthy subjects were placed on a WD for 1 month, the results showed a slightly decreased IL-6. Given that none of these clinical markers were significantly changed across all animals, this may suggest that these measures do not represent the underlying mechanisms changing plasma parameters after a short-term diet exposure. Similar to these plasma measures, the body weight of the five baboons changed differentially over the 49 d diet challenge and 57 d washout period (**Fig. 1).** At 49 d of WD exposure, three baboons gained weight, while two lost weight. During reversal to the monkey chow diet for 57 d, two baboons gained weight, two lost weight, and one maintained the weight gain status from the WD, suggesting high inter-individual variability in response to the short-term diet exposure. Interesting it has been shown in obese diabetic subjects with identical weight loss that the metabolite profiles are very different for gastric bypass surgery compared to a diet intervention [34]. It is likely that feeding behavior (frequency and amount) influences this, but these findings also suggest that the observed metabolic changes are likely not a result of weight gain but a primary response to WD exposure - not a secondary effect of weight gain.

**Fig 1.**
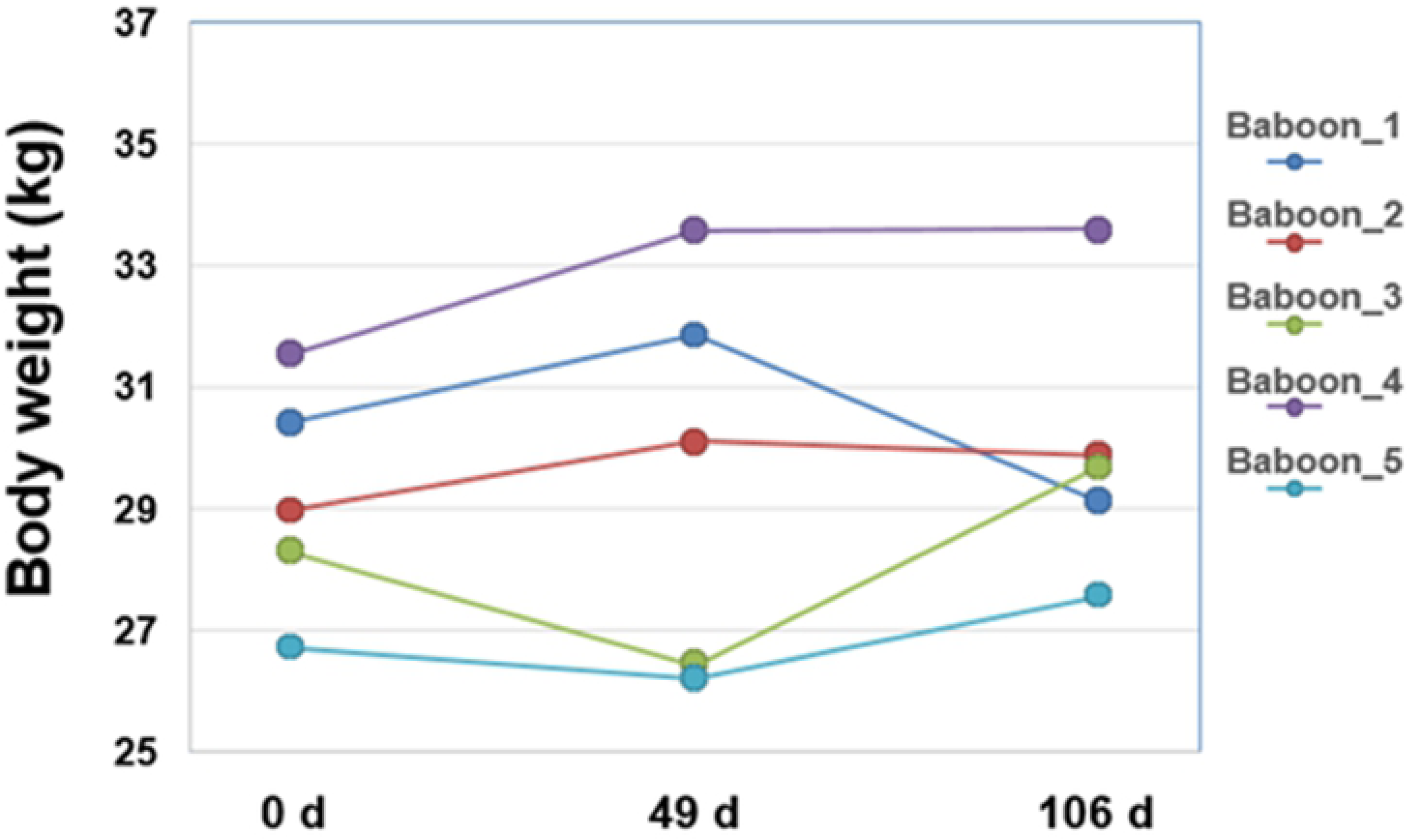
Variability in body weight reflecting individual response to diet challenge. Line chart showing change in body weight from 0 to 49 d and to 106 d for the five baboons in the study.

### 3.2. Response of clinical measures to the washout period after exposure to WD

Similar to the response to the WD, none of the changes seen in clinical measures after the washout period were statistically significant across all animals. We observed a few suggestive trends, including increases in insulin [2.94 fold at 106 d (767.26 ± 329.26 pg/ml) compared to 49 d (260.66 ± 88.41 pg/ml)], and circulating adipocytokines such as INF-gamma [4.11 fold increase at 106 d (33.64 ± 48.11 pg/ml) vs 49 d (8.17 ± 12.3 pg/ml)], IL-6 [12.15 fold at 106 d (9.11 ± 7.25 pg/ml) vs 49 d (0.75 ± 0.44 pg/ml)], TNF-alpha [2.44 fold at 106 d (2.5 ± 4 pg/ml) vs 49 d (1 ± 1.85 pg/ml)], and adropin [1.72 fold at 106 d (12.95 ± 8.1 ng/ml) vs 49 d (7.51 ± 9.9 ng/ml)] (**Table S1**). Comparison of these markers at 106 d to baseline (0 d) showed that concentrations did not return to baseline measures. These increased concentrations of clinical measures indicate that the 8 week chow diet after the WD challenge did not ‘washout’ the metabolic dysregulation induced by the WD, and suggests that the short-term exposure to the WD leads to long-term alterations in clinical plasma measures, suggesting systemic changes in metabolism in these animals. While none of these increases were statistically significant, the suggestive dysregulation of adipocytokine production may contribute to changes in insulin regulation and sensitivity, similar to the systemic insulin resistance reported in mice upon short-term HFD exposure, suggesting an induction of adipose tissue dysfunction [35]. However, inter-individual genetic variation likely influences these clinical measures, not just dietary intake, making them likely ill-suited as biomarkers to help monitor short-term responses in these animals.

### 3.3 Changes in baboon plasma metabolome in response to short-term WD exposure and upon return to chow diet

Metabolomics-based studies are rapidly maturing to allow the use of small molecule chemical profiling in complex biosystems research related to diet and nutrition [36]. Advances towards a clearer understanding of the role of specific dietary constituents in conferring cardioprotective and/or antidiabetic benefits require carefully controlled dietary intervention studies. Using a comprehensive 2D GC-MS platform, we obtained a snapshot of circulating metabolites in plasma reflecting the diet-induced physiological changes in baboons. We identified and quantified the relative amounts of 115 metabolites in the study at all-time points (**Table S2 and S3**). These metabolites covered 56 different KEGG-based metabolic pathways of which 21 were over-represented (P< 0.1), such as aminoacyl-tRNA biosynthesis, nitrogen metabolism, alanine, aspartate and glutamate metabolism, cyanoamino acid metabolism, glycine, serine and threonine metabolism, fatty acid biosynthesis, and D-Glutamine and D-glutamate metabolism among others (**Table S5)**.

Upon comparison of metabolite profiles between 0 and 49 d when WD was administered, we observed no statistically significant (P < 0.05) changes in individual metabolites. While we found trends for changes across the time course, similar to the clinical measures described above, due to the high variability (possibly from genotypic differences and thereby differential metabolic rates, and the small number of animals examined as part of our pilot study), there was no statistically consistent and significant difference between samples collected at 0 d and 49 d. However, we observed suggestive changes (P< 0.1) in 69 metabolites. Interestingly, all of these metabolites were ‘lower’ compared to the baseline (49 vs. 0 d; **Table S4)**. These changes included saturated fatty acids (caproic acid, capric acid, arachidic acid, lauric acid, butyric acid, pentanedecanoic acid) and other lipid compounds such as alpha-linoleic acid and glycerol (**Fig. 2).** Obesity and insulin resistance (IR) are associated with impaired elongation and desaturation of fatty acids, reflected in the serum by, e.g., higher proportions of myristic and palmitic acid and lower proportions of longer-chain n-6 and n-3 fatty acids [37, 38]. Decreased circulation of saturated fatty acids can also be considered beneficial because increased flux of saturated fatty acids into peripheral tissues can lead to accumulation of toxic lipid molecular species in these tissues [39, 40]. Plasma long-chain fatty acid concentrations have also been reported to be associated with the degree of obesity in humans [41]. Increased plasma glycerol concentrations are observed during the early phase of human obesity [42]. A human WD frequently contains excessive saturated and trans fatty acids [43].

**Fig 2.**
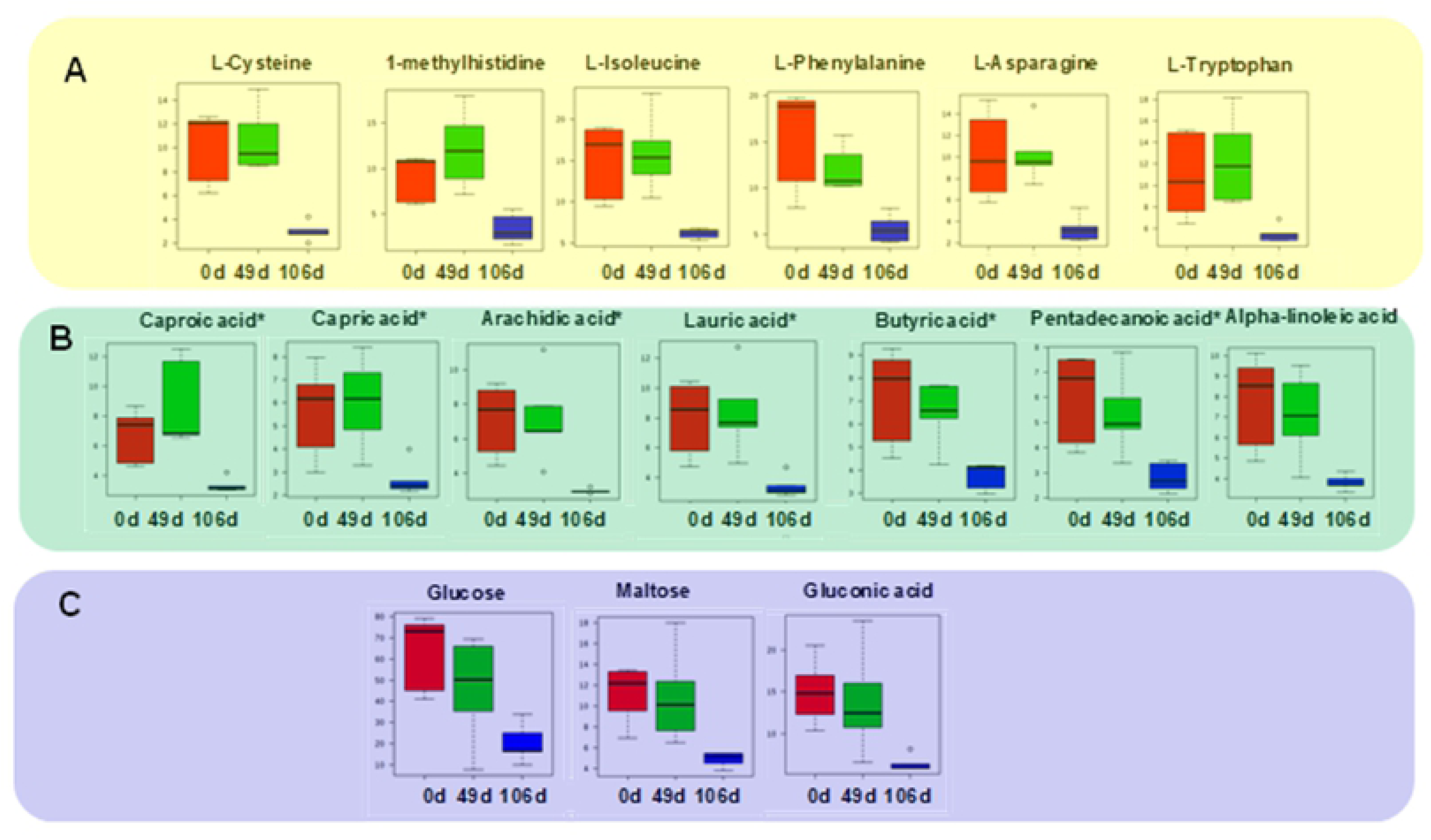
Relative abundances of different metabolites obtained from the metabolomics analysis. Box whisker plots (displaying the lower (Q1) and upper (Q3) quartiles and the median) showing relative metabolite abundance of **(A)** amino acids, **(B)** fatty acids (* are saturated fatty acids), and **(C)** carbohydrates, at each time point 0 d, 49 d, and 106 d. Only metabolites with P <0.1 are shown. P-values were obtained from within-subject ANOVA (for three time points 0, 49, and 106 d).

In addition to the lipid-related molecules, we observed changes in amino acids (1-methylhistidine, isoleucine, phenylalanine, asparagine, tryptophan, **Fig. 2**). Metabolic roles for amino acids are diverse, from providing substrates for mRNA translation, initiators of signal transduction and neurotransmission, biosynthesis of other nitrogen-containing compounds, to serving as substrates for gluconeogenesis, one-carbon methyl reactions, and tricarboxylic acid cycle [44]. The key role of amino acid metabolism early in the pathogenesis of diabetes and assessment of diabetes risk-management is well-established [45].

### 3.4. Clustering of diet-groups based on metabolite abundance

A major goal of this study was to compare the effects of the WD challenge and the washout period on quantity and quality of circulating plasma metabolites, as a function of systemic metabolic dysregulation. We observed that for time points 0 d, 49 d, and 106 d, two models explained the direction of changes for 105 of the 115 total metabolites we measured (**Fig. 3, Table S6**). The first model (P-value, 3.0 x 10^×27^, 85 metabolites) demonstrated a pattern of 0, 0, −1, indicating no changes to metabolites from time point 0 d to 49 d, followed by a decline in the levels of metabolites (−1) at 106 d. The 85 metabolites in the first model belonged mostly to amino acid metabolism (asparagine, glycine, aspartic acid, valine, isoleucine, leucine, threonine, tryptophan, tyrosine, proline, cysteine, and glutamic acid), nitrogen metabolism (glycine, hydroxylamine), butanoate metabolism (butanal, butyric acid, succinic acid), fatty acid biosynthesis (myristic acid, dodecanoic acid, and capric acid), cholesterol, maltose, myo-inositol, myo-inositol-phosphate, uridine, creatinine, canavanine among others. The second model (0, 1,-1) was based on 20 metabolites (P-value, 1.3×10^×7^) that belonged to amino acid metabolism (glutamine, serine, lysine, ornithine, 1-methyl histidine) ethanol, and glycerol.

**Fig 3.**
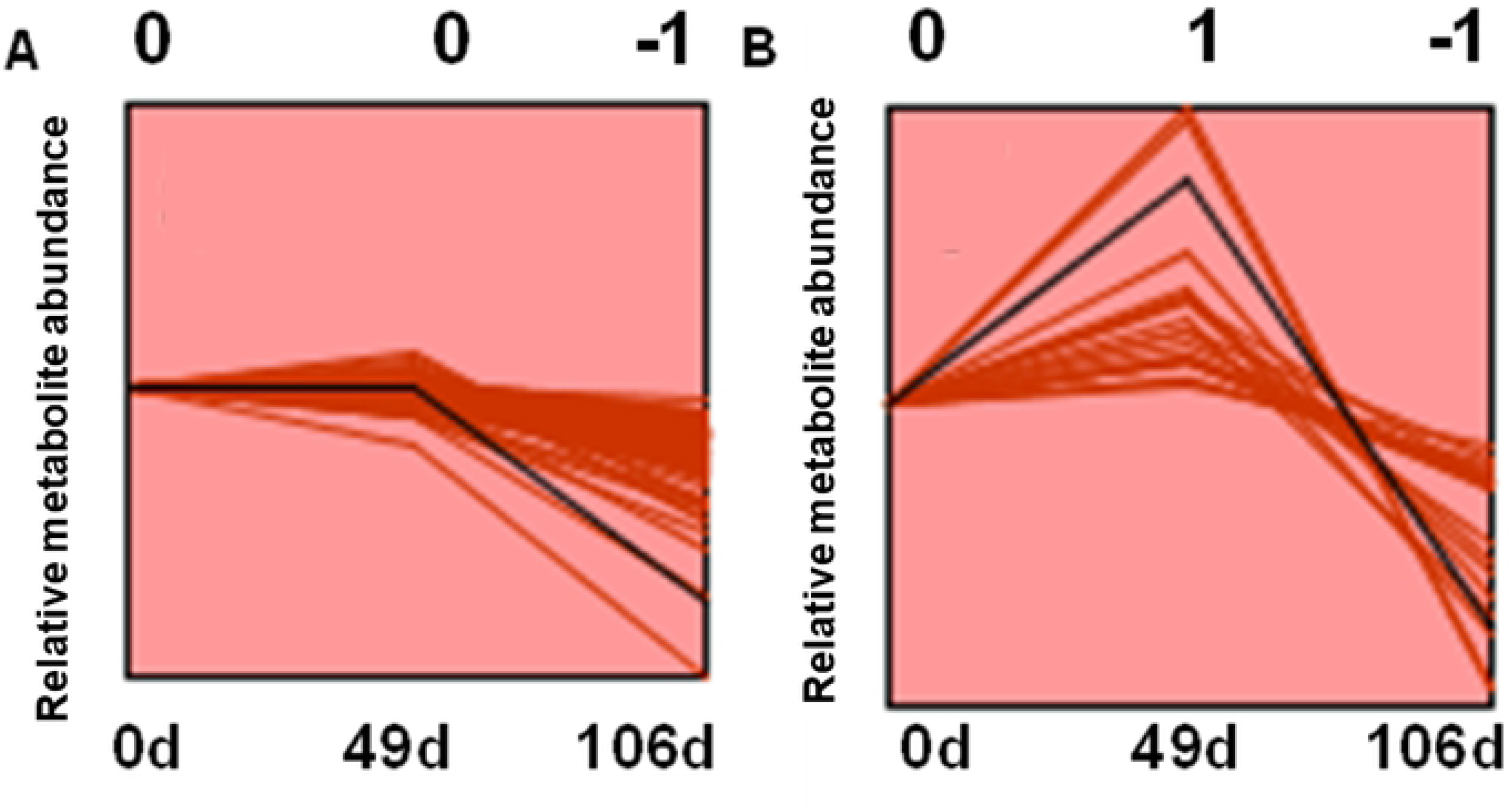
Patterns in changes of metabolite abundance at time pints 0 d, 49 d, and 106 d. For metabolites, the two models were significant with 85 and 20 metabolites showing patterns of 0 (unchanged), 0 (unchanged), −1 (lower) **(A)** and 0 (unchanged), 1 (higher), −1(lower) **(B)**, respectively.

It is striking that all metabolites in these two groups had lower circulating plasma levels at 106 d, when compared to the start of the study (0 d). Contrary to expectation, the levels did not simply return to baseline after washout, but settled on a different, lower “normal set point”, suggesting that the short-term WD challenge, regardless of the immediate impact during the diet administration, had long-term consequences, a process we have called ‘metabolic reprogramming” [46]. The most evident visualization of this phenomenon is provided in the plots in **Fig. 2** where levels are lower and variation among animals decreases at 106 d when compared to 0 d and 49 d.

To understand the contribution of the metabolite abundances in clustering and separation of the three time points of this diet challenge, hierarchical clustering analysis (HCA) was performed to classify the metabolites into clusters based on top 25 metabolites that were different for the group means (ANOVA). This clustering also suggests a tighter relatedness of 0 d with 49 d compared to metabolite levels after the washout period at 106 d (**Figure S1**). Moreover, the upper clusters were formed mostly by amino acids. Again, this analysis points to the clear metabolic differences in the 0 d and 106 d samples, confirming a “reprogramming” of the metabolic state due to the short-term exposure to the WD. This strongly suggests that the alterations in plasma metabolites after 106 d are truly reflective of a changes in underlying systemic metabolic regulation, and not simply a result of physiological changes such as weight gain induced during the WD exposure.

### 3.5. Metabolic dysregulation induced by WD prevented a return to baseline

We further looked into the metabolic changes in the baboons at 106 d as compared to the 0 d baseline levels. The dimension reduction strategy of unsupervised PCA was used to reduce all metabolites measured in plasma into a smaller number of orthogonal variables that revealed grouped and differential responses of the baboons to diet changes with an interpretable visualization. Upon PCA evaluation, the resulting plots explained ∼77 % of the variation by two PCs (PC1-61.8% and PC2-15.4%) attributed to the variation from the time (and diet) components (**Fig. 4A).** A supervised approach of classification, PLS-DA analysis, was conducted to indicate grouped responses of baboons to diet where the first two PCs explained 80% of the variations in the grouped responses (**Fig. 4B**). The clustering of 106 d samples reflects the reduced variability of the baboon metabolite abundances, and clearly separated from the 0 d and 49 d plasma samples.

**Fig 4.**
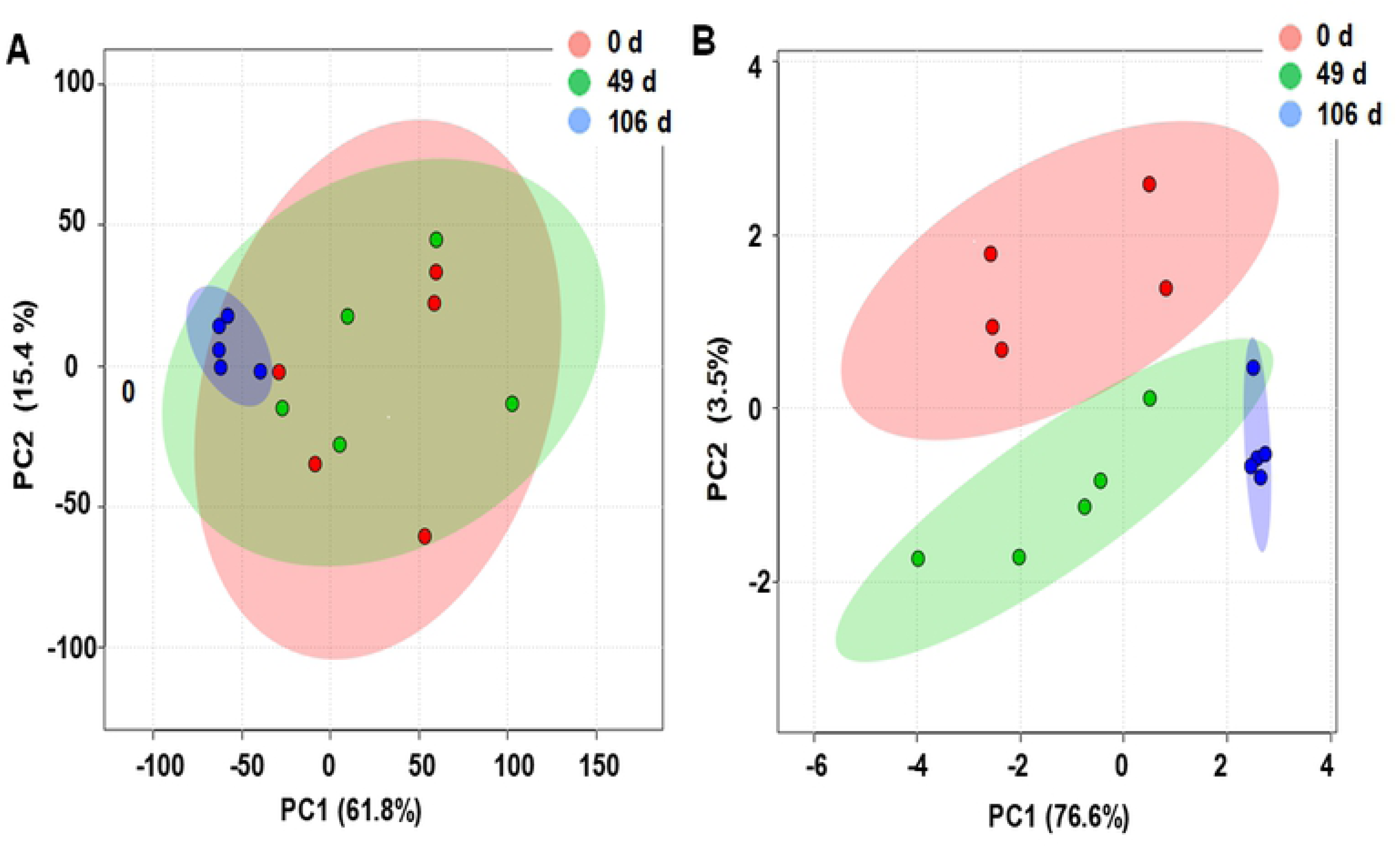
Multivariate analysis displaying score plots for supervised and unsupervised multivariate analysis. (**A**) PCA and (**B**) PLS-DA analysis. PCA was performed using five replicate data of relative metabolite abundances, and the generated PC1 and PC2 were plotted. PCA score plots display a clear separation of the sample groups (three time points) based on the abundance of 115 metabolites.

Further analysis of the metabolites driving the separation to identify the discriminatory metabolites using biplots derived from PLS-DA analysis revealed that sugars (glucose, galactose, mannose), urea, and fatty acids (palmitic and stearic acids) show higher discriminatory potential than others (**Fig. 5A**). Peining et al. [13] attributed this change to decreased catabolism due to increased calorie intake to explain changing metabolites at the time points in their weight loss study [13].

**Fig 5.**
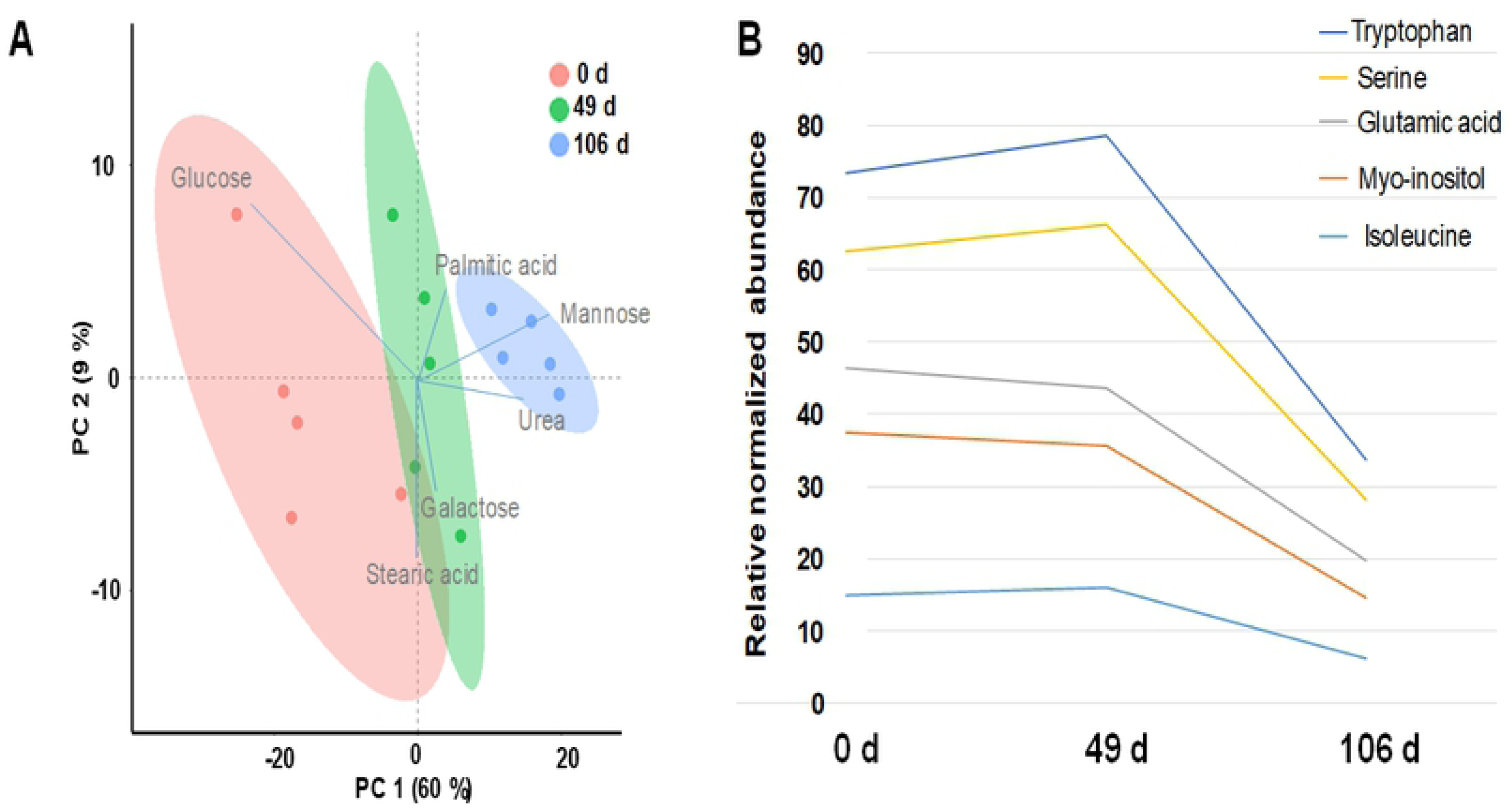
Metabolites contributing to the clustering of samples at the three time points. (**A**) A biplot from PLS-DA analysis displaying the metabolites contributing to the clusters of time points. For simplicity only the metabolites with higher discriminatory potential are shown. (**B**) A line chart displaying the trends of metabolites (amino acids and myo-inositol) over the study period.

Peining et al. also showed trends in metabolites such as isoleucine, myo-inositol, tryptophan, serine and glutamic acid [13] that changed in the same direction as the changes observed in our study. They reported a cluster of biomolecules that was increased upon weight gain but failed to return to baseline at the end of the weight loss period, including folate, phenylalanine and molecules related to branched chain amino acid (BCAA) degradation [13].

Even though several fatty acids did not show statistically significant changes, likely due to the small sample size (n=5) in this pilot study, the direction of the changes were consistent and qualitatively similar across these classes. In fact, different classes of lipids (for example, n-3 versus n-6) may have opposing net effects on specific processes, such as inflammation, and their relative proportions determine the net biological effect [47]. Most notably, measurements from this method are semi-quantitative, and reflect ion abundances for only a small subset of metabolites that are amenable to GC analysis, which may not always correlate with targeted methods that provide absolute concentration values. In addition, the statistical analysis of high-dimensional data is subject to overfitting and over interpretation because of the “small n/large p” paradigm. Despite these limitations of this pilot analysis, it is striking that the study clearly demonstrated that even after a relatively long washout period (diet with regular chow of 57 days), the metabolomic profiles do not return to pre-challenge levels, suggesting that the diet exposure has long-term metabolic consequences on the animals, independent of weight gain (the study design reported elsewhere [13]) or any other consistent clinical or pathological changes.

While the changes in plasma metabolites observed in our analysis suggest an interesting long-term physiological response to the 7 week WD challenge in our animals, further studies will be needed to elucidate mechanisms and address weaknesses in our existing pilot. Most notably, the findings will need to be confirmed in a larger number of animals. Finally, future studies may require collection of food consumption rates for individual animals, a factor that may have influenced our pilot analysis, and may allow a more detailed analysis of genetic factors contributing to the variability in individual responses. Therefore, we consider the modeling results presented here as hypothesis generating until confirmed with more targeted approaches that allow a more comprehensive and accurate correlation of genetic predisposition, systemic physiology, and plasma metabolite changes. Nonetheless, despite the underlying inter-individual variability and the small sample size, the consistent finding of altered plasma metabolite concentrations even after a return to a normal diet, and the reduced variability observed across the animals suggests a systemic regulatory mechanism that alters these metabolites in response to a WD diet in a tightly controlled way. This metabolic reprogramming may provide a first insight into potential pathophysiological mechanisms contributing to the negative health consequences of weight cycling and short-term diet changes in humans.

## 4. Concluding remarks

In our baboon study, short-term exposure to a WD did not induce significant metabolomic changes after 7 weeks (49 d). However, samples collected after the washout period of 57 d showed large-scale metabolic changes in these animals, resulting in metabolite levels that were lower relative to the 0 d time point, and less variable across all animals. The post-washout period metabolite levels did not revert back to the 0 d baseline abundance levels, suggesting metabolic reprogramming with the establishment of a tightly regulated and controlled new “normal set point” in response to the short-term WD challenge. These results provide new insights into potential targets and related pathways that reflect the metabolic health status in baboons. Some of these metabolites and metabolic signatures may be indicative of long-term physiological changes and chronic disease susceptibility. Additional studies will be required to further elucidate the metabolic consequences of these changes in plasma metabolites, and how this alters the metabolic regulation in these animals.

## Author Contributions

AC, LAC, MO designed the research; VM, RPU, BBM conducted the research; MO, LAC, AC provided essential reagents and materials; BBM analyzed and interpreted data, BBM, MO, LAC, JSP wrote the manuscript; and have primary responsibility for the final content and edits. All authors read and approved the final manuscript.

## Disclosures

None.

### Acknowledgements

Research reported in this manuscript was supported by a grant in aid from the National Institutes of Health (NIH): P01 HL028972. This investigation used resources, which were supported by the Southwest National Primate Research Center grant P51 RR013986 from the National Center for Research Resources, NIH, currently supported by the Office of Research Infrastructure Programs (ORIP), NIH through grant P51 OD011133. This investigation was conducted in facilities constructed with support from ORIP through grant numbers C06 RR14578, C06 RR15456, C06 RR013556, and C06 RR017515.

## Supporting Information

**S1 Table. Clinical measures (raw data) obtained from the 5 animals fed WD diet for 49 d and brought back to chow diet.**

**S2 Table. List of 115 metabolites identified and their raw relative abundances obtaining using the 2D GC-ToF-MS platform alongside their identifiers.** Identifiers are KEGG: Kyoto Encyclopedia of Genes and Genomes, HMDB: Human Metabolome Database, PubChem ID, ChEBI, METLIN IDs.

**S3 Table. Processed, normalized and transformed relative metabolite abundance data for the three time points, 0, 49, and 106 d as well as the mean, SD, RSD, and fold changes (FC).**

**S4 Table. ANOVA and post-hoc (Fisher’s LSD) results for the metabolite abundance data conducted on the three time points.**

**S5 Table**. **Enrichment analysis for KEGG-based pathways represented by the metabolites quantified in this study.**

**S6 Table. Pattern analysis displaying the list and values of 85 and 20 metabolites fitting into two models, (0, 0, −1) and (0, 1, −1), respectively.**

**S1 Fig. Hierarchical cluster analysis (HCA) of mean and normalized abundances of metabolites obtained from five biological replicates across the time points 0, 39, and 106 d. Red and green indicate increasing and decreasing concentrations of metabolites, respectively.** Values were subjected to average linkage clustering (Pearson distance). Scale indicates high (3, red) to green (−3, low) abundances.

## List of abbreviations

2D GC-ToF-MS: Two dimensional gas chromatography time-of-flight mass spectrometry,
ANOVA: analysis of variance,
BCAA: branched chain amino acid,
CVD: cardiovascular diseases,
HCA: hierarchical clustering analysis,
HDL: high density lipoprotein,
KEGG: Kyoto Encyclopedia of Genes and Genomes,
LDL: low density lipoprotein,
*m/z*: mass-to-charge ratio,
MeOX: methoxyamine hydrochloride,
MSTFA: N-methyl-N-trimethylsilyl-trifluoroacetamide,
NHP: non-human primate,
PCA: principal component analysis,
PUFA: polyunsaturated fatty acid,
QC: quality control,
WD: Western diet,

